# Optimized breath detection algorithm in Electrical Impedance Tomography

**DOI:** 10.1101/270348

**Authors:** D. Khodadad, S. Nordebo, B. Müller, A. Waldmann, R. Yerworth, T. Becher, I. Frerichs, L. Sophocleous, A. van Kaam, M. Miedema, N. Seifnaraghi, R. Bayford

## Abstract

**Objective:** This paper define a method for optimizing the breath delineation algorithms used in Electrical Impedance Tomography (EIT). In lung EIT the identification of the breath phases is central for generating tidal impedance variation images, subsequent data analysis and clinical evaluation. The optimisation of these algorithms is particularly important in neonatal care since the existing breath detectors developed for adults may give insufficient reliability in neonates due to their very irregular breathing pattern.

**Approach:** Our approach is generic in the sense that it relies on the definition of a gold standard and the associated definition of detector sensitivity and specificity, an optimisation criterion and a set of detector parameters to be investigated. The gold standard has been defined by 11 clinicians with previous experience with EIT and the performance of our approach is described and validated using a neonatal EIT dataset acquired within the EU-funded CRADL project.

**Main results:** Three different algorithms are proposed that are improving the breath detector performance by adding conditions on 1) maximum tidal breath rate obtained from zero-crossings of the EIT breathing signal, 2) minimum tidal impedance amplitude and 3) minimum tidal breath rate obtained from Time-Frequency (TF) analysis.

**Significance:** Based on the gold standard, the most crucial parameters of the proposed algorithms are optimised by using a simple exhaustive search and a weighted metric defined in connection with the Receiver Operating Characterics (ROC). This provides a practical way to achieve any desirable trade-off between the sensitivity and the specificity of the detectors.

## 1. Introduction

Robust and reliable detection and delineation of breaths known as the breath detection is the basis of lung functionality analysis. The aim of the breath detector is to obtain information about the exact timing of the two respiration phases: inspiration and expiration. Accurate breath detection is a key parameter in the clinical assessment and monitoring of respiratory function. However, this process is technically challenging due to the noisy nature of the acquired data obtained from the thorax area, measurement artifacts and other factors such as the presence of sighs, swallows, transient reductions (hypopnoeas and bradypnoeas) and pauses (apnoeas) in breathing. In order to quantify key breathing parameters, accurate identification of inspiration and expiration phases in each breath cycle is required.

Different methods to quantify respiration have been attempted during the last years. Measuring airflow and the derived volume signal, is an example of the traditional method which need a mouthpiece, a breathing mask or an endotracheal tube [1, 2]. For example, airflow curves are used as the basic delineator while the airway pressure and the CO_2_ concentration curves are used to confirm the delineation [3]. However, the method is time consuming and potentially prone to human error and indeed impractical for large data sets. In order to cope with these problems, automated physiological landmark detection in the airflow or epiglottis pressure signal (Pepi) methods are introduced [4]. However, the presence of baseline volume shifts, can still render these algorithms inaccurate.

Similar to the breath airflow methods, breath sound detection algorithms are also developed to detect the breath cycle. Although these methods are used in clinical practice, they are mostly developed for industrial applications such as breath sound removal/suppression in the music industry and monitoring firefighters’ respiration during their duties. These methods are based on a template matching approach, and therefore, they are more reliable for adult normal breath detection [5]. Recently, transcutaneous electromyography of the diaphragm (dEMG) is used for non-invasive breath detection based on the measurement of electrical activity of the diaphragm [6, 7].

Chest electrical impedance tomography (EIT) is a non-invasive monitoring bedside tool for imaging regional electrical impedance or voltage changes induced by changes in the regional lung ventilation. Computed Tomography (CT) and Magnetic Resonance Imaging (MRI) are established medical methods for imaging lung anatomy, however, they cannot be used at the bedside and do not allow the assessment of lung function. Online X-ray monitoring is not possible because it would increase the radiation load due to the need for successive imaging. Further, these modalities are incompatible with the neonatal application as they demand full patient corporation to be stable, which is not feasible [7–10]. Furthermore, CT and MRI do not produce dynamic images and it is not possible to achieve continuous monitoring of regional lung ventilation in Neonatal Intensive Care Unit (NICU). Unlike CT or other radiographic techniques, EIT is a relatively inexpensive technology which makes continuous real-time radiation-free monitoring of the lung function possible directly at the bedside without any known hazard. The EIT therefore can be an ideal candidate to be applied for monitoring neonates with mechanically supported ventilation [8–14].

The main scope of this study is to define a specific clinical parameter using an EIT device to achieve an optimised breath delineation for neonates and premature newborns that may have irregular breathing pattern [15]. A significant number of studies for lung function monitoring using EIT have been published in recent years [16–20]. There are different methods to resolve the respiratory related information from the chest EIT signal. Peak and slope detection based on the breathing (impedance) signal is straightforward in order to determine the breathing cycles in tidal breathing [4, 5, 21]. Indeed, the slope of the breathing signal shows whether the global impedance is increasing or decreasing in order to determine the inspiration and expiration phases. However, due to the presence of cardiac related signal components and other disturbances, the method is prone to yield high error rates. Frequency domain filtering is attempted as a solution to decompose and extract cardiac and respiratory related signals [17, 22, 23]. However, the method often suffers from a frequency overlap of the respiration harmonics and the heartbeat-related signal. In order to achieve a proper separation of the respiratory and cardiac-related signals, Principal Component Analysis (PCA) may be used. The PCA demands identification of template functions for time domain filtering. It is a multivariate statistical method and also time consuming which makes the method not suitable for real time analysis [18]. Consequently, Independent Component Analysis (ICA) has been used to increase the accuracy of decomposing the EIT signal into independent components based on statistical characteristics of the signals [24].

Three different methods to obtain breathing index from EIT are evaluated:

1. the zero-crossing (ZC) algorithm,
2. the zero-crossing algorithm with amplitude threshold (ZC-AT),
3. the zero-crossing algorithm with amplitude threshold and FFT-based breath rate estimation (ZC-AT-FFT).

The algorithms use the global impedance signal obtained from EIT defined as the sum of all pixel values and are improving the detector performance by adding conditions on 1) maximum tidal breath rate obtained from zero-crossings of the EIT breathing signal, 2) minimum tidal amplitude and 3) minimum tidal breath rate obtained from Time-Frequency (TF) analysis (apnea alarm). In analogy to [5, 21] we determine the breathing cycle from the position of peaks and slopes in the breathing signal and add an extra criterion regarding the minimum distance in between zero crossings at one half and a full cycle, respectively. In this way, higher oscillating error sources are avoided. In the second algorithm, an amplitude threshold is used to remove small amplitude fluctuations (e.g., cardiac related) which are not breaths. Finally, in the third algorithm, the Short-Time Fourier Transform (STFT) is employed to dynamically estimate the breath rate, facilitating a lower bound on what is realistically considered to be a breath (e.g., by detecting hypopneas and apneas) [25]. This is an advantage as to our knowledge, up to now no EIT device shows respiratory rate, based on the EIT measurements.

The most crucial parameters of these algorithms are optimised by using a simple exhaustive search and a weighted metric defined in connection with the Receiver Operating Characterics (ROC). This provides a practical way to achieve any desirable trade-off between the sensitivity and the specificity of the detectors. The approach is generic and is optimised and validated using a dataset with clinical neonatal EIT data from an ongoing study within the EU-funded project <u>CRADL</u> (Continuous Regional Analysis Device for neonate Lung).

The gold standard is defined based on the training data examined by 11 clinicians with previous experience with EIT. It is emphasised that this approach is non-parametric in the sense that no statistical assumptions have been made on the population distributions from which the data are drawn. To this end, the non-parametric gold standard constitutes a generic approach which is currently under development and is expected to be useful in many other EIT related issues and investigations where breath delineation and tidal breathing is of great importance, see e.g., [26].

## 2. Methods

### 2.1. Data acquisition

Data collection is performed within an ongoing prospective clinical study as part of the CRADL project. Infants with a body weight less than 600 g, postmenstrual age less than 25 weeks at inclusion, electrically active implants or those suffering from thorax skin lesions were excluded from the study. Informed written consent was obtained from the parents of the neonatal study participants. The raw EIT data were acquired by the CRADL study EIT device (Swisstom, AG, Landquart) with 32 electrodes at the frame rate of 48 Hz. This system is specially designed for infants who had thorax diameters as small as 17.5 cm [27]. Current injections with amplitude of 3 mA rms at a frequency of 200 kHz were applied using a skip 4 injection pattern. The resulting voltage differences were measured by the remaining electrode pairs after each current injection. EIT data were filtered using a bandpass filter with cut-off frequencies of 0.15 Hz and 1.8 Hz in order to remove DC and cardiac related impedance changes. The GREIT reconstruction algorithm [20] was then used to reconstruct the images.

### 2.2. Breath detection

In the following, the description of the three proposed algorithms and their related parameters are given. Subsequently, the definition of the gold standard, the optimisation and the validation of the methods are presented.

#### 2.2.1 Zero-crossing algorithm

The zero-crossing (ZC) algorithm constantly monitors the time instances of zero crossings in the breath impedance signal and operates with the two states “Wait for Raising Crossing” (WRC) and “Wait for Falling Crossing” (WFC), see Figure 1. The “Identical Crossing Spacings” (ICS) and “Different Crossing Spacings” (DCS) in the breath signal are defined as the differences in the time of consecutive zero crossings with identical or different slopes, respectively. To mitigate the detection of rapid oscillations as being breaths, two threshold parameters are defined, the Minimum Identical Crossing Spacing (MICS) and the Minimum Different Crossing Spacing (MDCS) in seconds. These parameters are also normalised to the maximal breath period so that

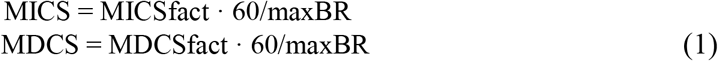

where MICSfact and MDCSfact are the normalised parameters and maxBR the assumed maximal breath rate in breaths/min.

**Figure 1.**
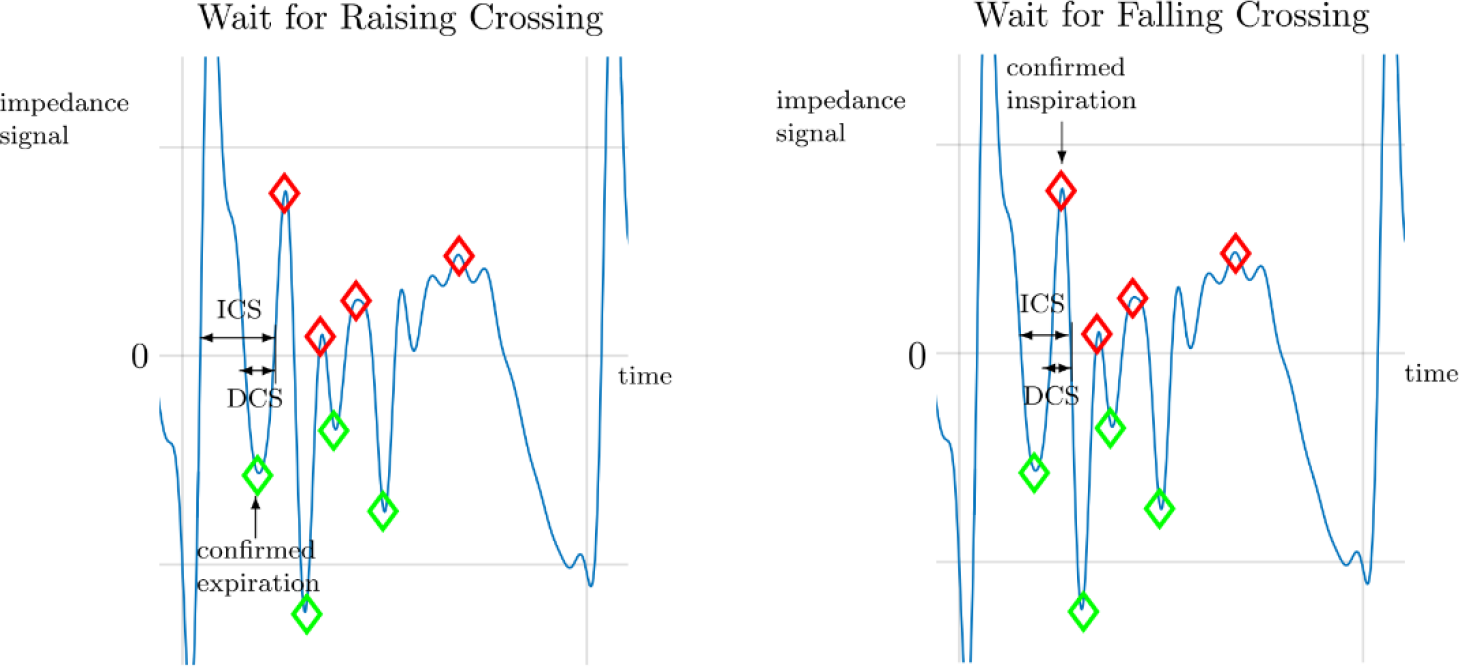
Illustration of the zero-crossing algorithm with Identical Crossing Spacing (ICS) and Different Crossing Spacing (DCS) indicated in the breath (impedance) signal. The green and red diamonds indicate the detected minima and maxima during the expiration and inspiration phases, respectively. Left: end of WRC state and confirmed expiration phase. Right: end of WFC state and confirmed inspiration phase.

During the WRC state, the algorithm is constantly updating the current minimum of the signal. If a raising zero crossing is detected and both criteria ICS > MICS and DCS > MDCS are satisfied, then the last minimum is confirmed as the last expiration phase and the state of the algorithm is changed to WFC. Similarly, during the WFC state, the algorithm is constantly updating the current maximum of the signal. If a falling zero crossing is detected and both criteria ICS > MICS and DCS > MDCS are satisfied, then the last maximum is confirmed as the last inspiration phase and the state of the algorithm is changed to WRC.

Tidal amplitudes are defined as the differences in signal impedance amplitudes between the maxima during the inspiration phases and the corresponding minima during the expiration phases. Here, it is assumed that the expiration phase precedes the inspiration phase during one breathing cycle.

With the ZC algorithm, the maximal breath rate parameter is set to maxBR = 150 breaths/min and the parameters MICSfact and MDCSfact are optimised as described below.

#### 2.2.2 Zero-crossing algorithm with amplitude threshold

To decrease the rate of False Positives (FP) in breath detections due to superimposed small amplitude changes, such as cardiac related impulses during short periods of no breathing or other disturbances, we propose using an amplitude threshold which has proven to be very efficient. A statistical based threshold is therefore used here as follows. It is assumed that a finite record of data is going to be analyzed where the majority of signal samples (at least more than, e.g., 50%) come from the measurement periods containing breathing signals and not only noise.

To determine a typical value representing large tidal amplitudes, we choose the tidal amplitude typTA at a certain upper percentile (UP) of all tidal amplitudes in the record. This is to avoid comparing to some large impulsive disturbances, noise or other signal artifacts during the recording. In this study the value UP = 0.8 has emerged as a good compromise during initial testing and has been fixed during the optimisation. A lower threshold for tidal amplitudes is then determined as lowTA = typTA·lowTAfact where lowTAfact is a lower amplitude factor chosen in the range 0 < lowTAfact < 1. In the breath detection algorithm, whenever a tidal amplitude detected by the zero crossing algorithm has a tidal amplitude lower than lowTA, it will be discarded as a tidal amplitude and not counted as a breath. With the ZC-AT algorithm, the maximal breath rate parameter is set to maxBR=150 breaths/min, the upper percentile UP=0.8 and the parameters MICSfact, MDCSfact and lowTAfact are optimised as described below.

#### 2.2.3 Zero-crossing algorithm with amplitude threshold and FFT-based breath rate estimation

A Short-Time Fourier Transform (STFT) is implement on the breath impedance signal *x(n)* which is given by the sum of pixel (impedance) values obtained from EIT - see details in [20]. The STFT is given here by the following Fast Fourier Transform (FFT) calculation [28] at discrete frequency and time *(k, n)*

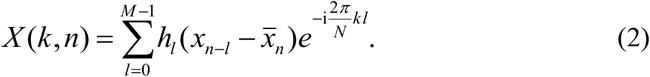

Here, a sliding time window is employed where *x*_*n-l*_ denotes the finite sequence of temporary data to be analyzed, *h*_*l*_ the corresponding window weight function of length *M, N* the size of the FFT, and where the discrete impedance signal *x(n)* has been sampled at the frame rate *f_s_*. The mean 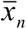 is calculated as 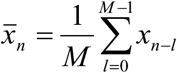.

A crucial step of any STFT implementation is to determine a suitable trade-off between the resolution in time and frequency. Based on the available set of patient data, we have found that a suitable time window for analysis in our application is about 3 seconds in duration yielding a window length of 143 samples at a frame rate of 48 Hz. A zero padding is used with the parameter *N* = 1024 yielding a FFT frequency resolution of 2.8 breaths/min. A Kaiser window [28–30] is employed to achieve an optimal trade-off between the width of the main-lobe (actual frequency resolution) and the side-lobe rejection in the frequency domain. Here, the Kaiser window was chosen with a main-lobe width of about 30 breaths/min and a side-lobe rejection better than 30 dB. Finally, the breath rate estimate BR(*n*) is obtained as the breath rate for which the STFT |*X*(*k,n*)| has its maximum at time *n*, see Figure 2.

**Figure 2.**
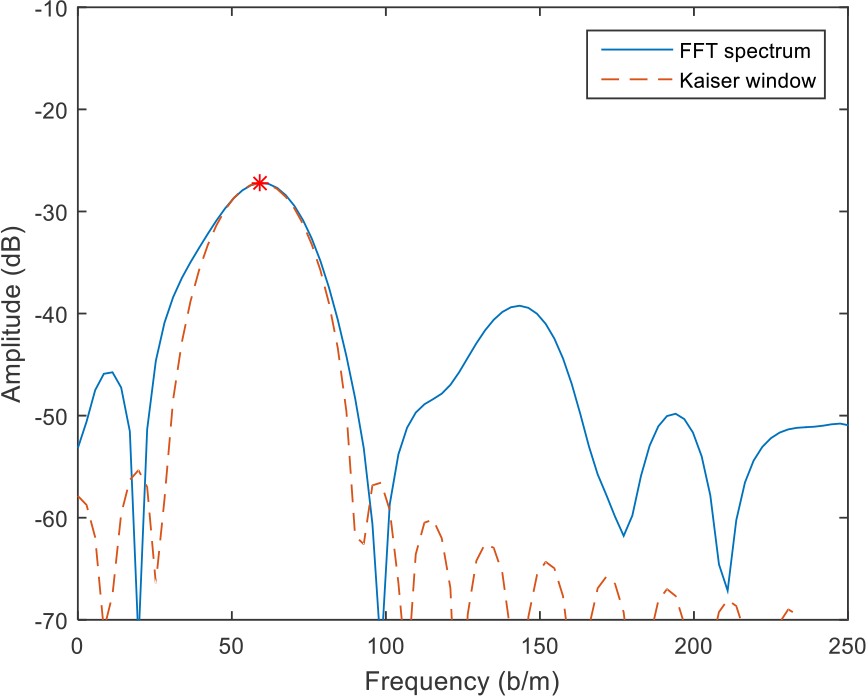
STFT spectrum of the EIT data (blue line) and Kaiser window (dashed line). The respiratory rate estimate is defined as the frequency for which the STFT has its maximum (red star).

Note that in (2), it is necessary to subtract the mean 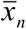 of the temporary data vector to avoid spectral leakage from the zero-frequency component (DC level) in the FFT. However, further improvement is also obtained by pre-processing the data using a digital high-pass filter to remove the dominating low frequency contents. Here, a second order high-pass Butterworth filter has been used with a cut-off frequency of 15 breaths/min.

A breath alarm (BA) level is defined in breaths/min. In the breath detection algorithm, whenever a tidal amplitude detected by the zero-crossing algorithm has a tidal amplitude lower than lowTA, or the estimated breath rate BR(n) is lower than the alarm level BA, it will be discarded as a tidal amplitude and not counted as a breath.

With the ZC-AT-FFT algorithm, the maximal breath rate parameter maxBR is dynamically set to maxBR(*n*) = 4·BR(*n*) breaths/min, the upper percentile UP = 0.8 and the parameters MICSfact, MDCSfact, lowTAfact and BA are optimised as described below.

### 2.3 Optimisation and validation of breath detection

In order to perform an optimisation and validation of the breath detection algorithms, there is a need for a gold standard. In the following, details of defining the gold standard and the procedure of optimisation and validation will be described.

#### 2.3.1 Gold standard for breath detection

To optimise and validate the breath detection algorithms two sets of data were selected, a test set and a validation set, each set comprising 10 records of CRADL patients’ data (impedance signals) with a duration of 80 seconds each.

11 experienced neonatologists working with EIT, were asked to examine the corresponding impedance plots and to indicate which peaks correspond to a regular breath cycle, and ignore other peaks. The 80% limit of agreement was defined as a gold standard. It was therefore decided to define as a true breath (or positive breath) all peaks that at least 80% of the clinicians agreed it was a breath, and to mark the other peaks as no breath (or negative breath). The result of the examination (gold standard for breath detection) is illustrated in the Figures 5-6 showing an example of the breath impedance signals together with a red and a black asterisk for a positive and a negative breath, respectively.

#### 2.3.2 Optimisation

The test data sets have been used for training in order to optimise the breath detection algorithms ZC, ZC-AT and ZC-AT-FFT, in terms of their Receiver Operating Characteristics (ROC) defined by the corresponding true positive rate (TPR) and false positive rate (FPR) [31]. The validation data sets have then been used to validate the results in terms of the ROC. The number of True Positives (TP), False Negatives (FN), False Positives (FP) and True Negatives (TN) are readily calculated based on the output of each detector in comparison to the gold standard. The TPR and FPR are then calculated based on all the 10 data sets in either the test set or the validation set, respectively.

To define a practical optimisation problem in view of the multi-criteria ROC plot, we employ the weighted norm (a weighted metric distance from the optimal point (FPR, TPR) = (0, 1))

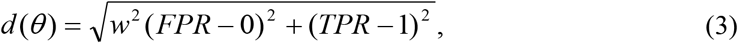

where *θ* is the parameter vector (e.g., *θ* = (MICSfact, MDCSfact, lowTAfact, BA) for the ZC-AT-FFT algorithm), FPR and TPR are the false and true positive rates obtained with the current parameters, and *w* is a positive weighting factor. The purpose of the weighting is to parameterise the trade-off between the conflicting requirements to have a large TPR (high sensitivity) and a small FPR (high specificity) at the same time. In particular, in our example, we are mostly concerned with emphasizing a small FPR, and a suitable weight for this purpose was found to be *w* = 10.

The discrete nature of this global optimisation problem makes the problem very complex. In particular, the parameters of the algorithms are not independent and the change of one parameter may be compensated by a change of another parameter making the solution non-unique. Since the corresponding global optimisation problem becomes huge when increasing the number of degrees of freedom, a practical approach is to use the optimisation software to make experiments and to find a good compromise regarding which parameters realistically can be fixed and which parameters need to be optimised. In this sense, the resulting parameters will always be suboptimal. The results presented below constitute one such compromise and serve the purpose to illustrate the proposed optimisation methods and to quantify the (suboptimal) performances of the breath detectors that have been studied.

To this end, the most important parameter of the detectors ZC-AT and ZC-AT-FFT is the lower amplitude factor lowTAfact for which it was easy to find an optimal interior solution with 0 < lowTAfact < 1.

## 3. Results

As explained in section 2, the proposed metric distance function (3) with a weight *w* = 10 is used to optimise ZC, ZC-AT and ZC-AT-FFT algorithms. Figure 3 shows an example of choosing the optimal parameters based on defining a minimum distance from the perfect optimal point (TPR = 1, FPR = 0). The corresponding metric distances for each possible choice of the parameters are calculated, and shown (sorted) in Figure 3. Figure 4 shows a typical example of determining an optimal lowTAfact when the other parameters are fixed on their optimal values. The search for the optimal lowTAfact has been done in the range of [0 0.3] with the step forward of 0.001 when MICSfact, MDCSfact and BA are fixed on 0.3, 0.1 and 30, respectively. The most important optimisation parameter is lowTAfact where an interior solution satisfying strict bounds 0<lowTAfact<1 could readily be found. With the other parameters, due to their inherent redundancy, the optimal solution always saturated at the given upper or lower parameter bounds. Hence, considering the discrete nature of this global optimisation problem, a relatively small parameter space has been used together with an exhaustive search for minimizing the metric distance function (3) with weight *w* = 10. The following results have been obtained:

- Zero-crossing (ZC) algorithm.

Default parameters: MICSfact=0.75 and MDCSfact=0.25.
Validated performance: FPR=0.23, TPR=0.96.
- Zero-crossing (ZC) algorithm.

Parameter range: 0.5 ≤ MICSfact ≤ 0.75 and 0.1 ≤ MDCSfact ≤ 0.25.
Optimised parameters: MICSfact=0.5 and MDCSfact=0.1.
Validated performance: FPR=0.23, TPR=0.97.
- Zero-crossing algorithm with amplitude threshold (ZC-AT).

Parameter range: 0.5 ≤ MICSfact ≤ 0.75, 0.1 ≤ MDCSfact ≤ 0.25 and 0.1 ≤ lowTAfact ≤ 0.3.
Optimal parameters: MICSfact=0.5, MDCSfact=0.1 and lowTAfact = 0.25.
Validated performance: FPR=0.08, TPR=0.95.
- Zero-crossing algorithm with amplitude threshold and FFT-based breath-rate estimation (ZC-TA-FFT).

Parameter range: 0.5 ≤ MICSfact ≤ 0.75, 0.1 ≤ MDCSfact ≤ 0.25,
0.1 ≤ lowTAfact ≤ 0.3 and 0 ≤ BA ≤ 40.
Optimal parameters: MICSfact=0.5, MDCSfact=0.25, lowTAfact = 0.15 and BA = 30.
Validated performance: FPR=0.06, TPR= 0.84.

**Figure 3.**
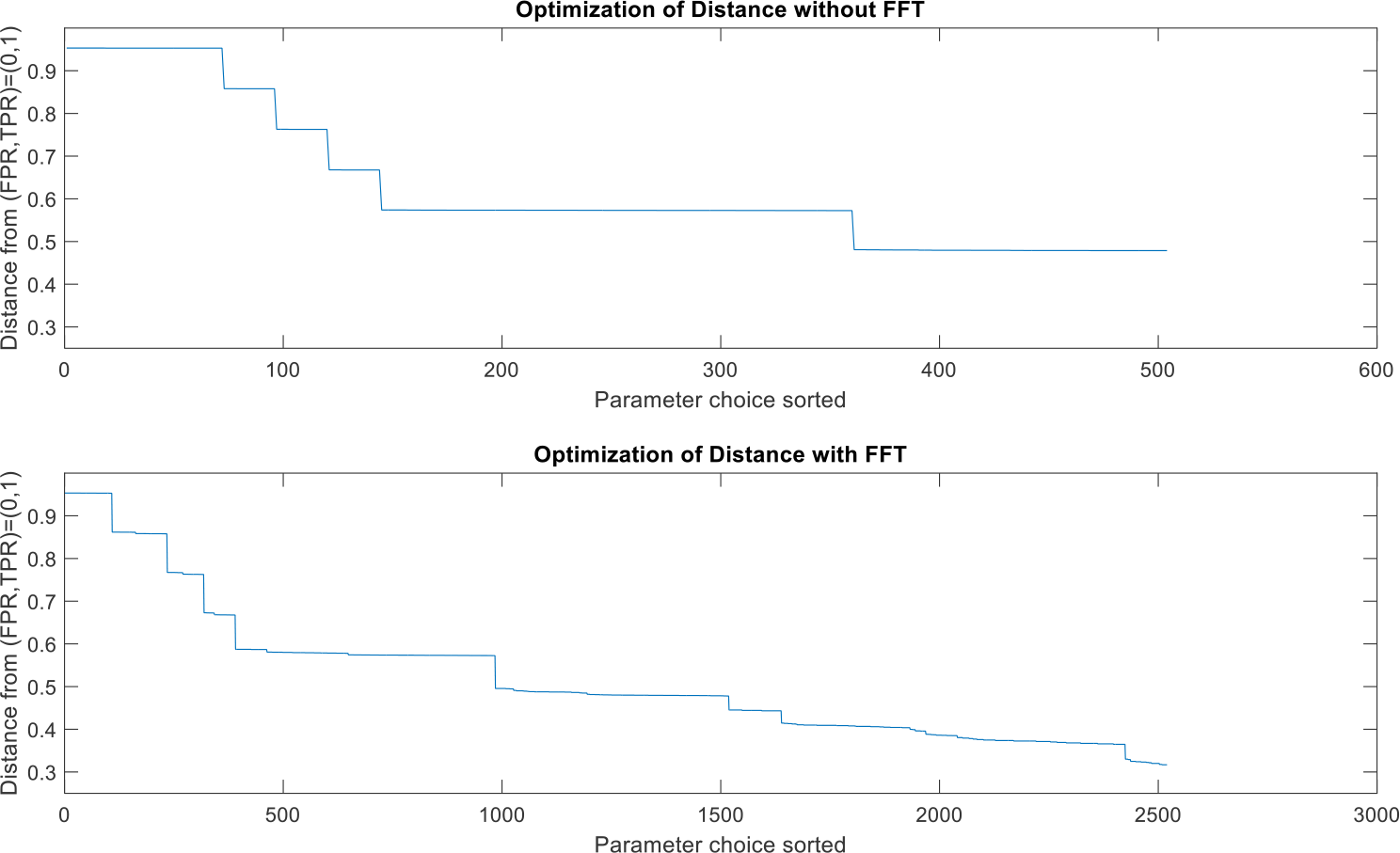
Weighted metrics over the chosen parameter set in the optimisation of the ZC-AT and the ZC-AT-FFT algorithms, respectively.

**Figure 4.**
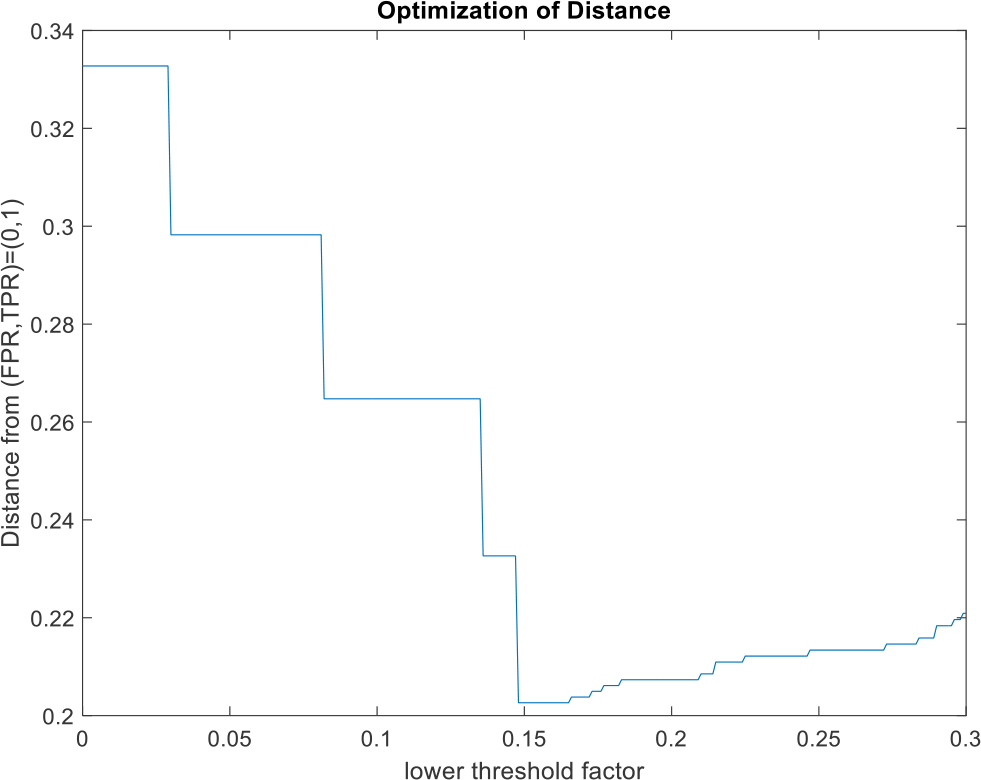
A typical example of determining an optimal lowTAfact when the other parameters are fixed on their optimal values.

Figure 5 shows a result of using the ZC-AT algorithm with optimal parameters for a typical breath detection example using the validation data set. The upper plot shows the breath impedance signals together with a red and a black asterisk for a positive and a negative breath, respectively. Four-second intervals at the beginning and at the end of each data have been excluded from the investigation, allowing the breath detectors to stabilise and avoid edge effects. The lower plot shows the output of the optimised ZC-AT detector. The performance in terms of TP, FN, FP and TN is shown in the title of the upper plot. It is seen that the ZC-AT algorithm performs with 36 TPs and 3 FPs (indicated in the figure).

**Figure 5.**
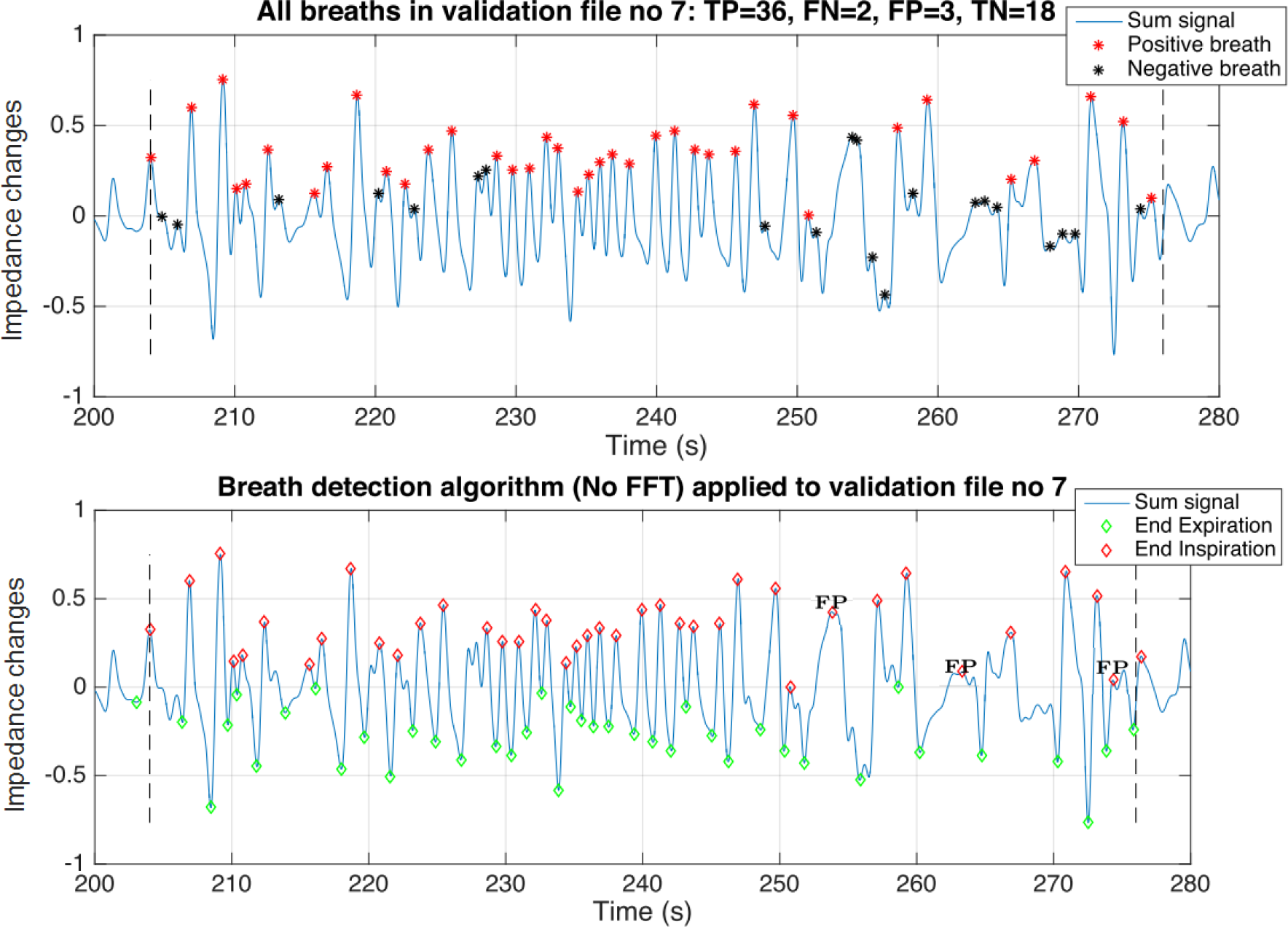
Breath detection performance on the validation data no.7 with zero-crossing algorithm complemented with amplitude threshold (ZC-AT). FP represents false positive.

In Figure 6 the ZC-AT-FFT algorithm with optimal parameters is used for the same particular breaths as in Figure 5. The second and third plots show the output of the optimised detector using the ZC-AT-FFT algorithm and the corresponding estimated breath rate, respectively. The detector performs with 1 FP and 29 TPs for the same data. It is easy to identify the particular breaths where the ZC-AT-FFT algorithm is able to avoid 2 of the FPs by employing a breath alarm level (BA). However, this is done with the cost of neglecting TPs in the area/situations with breath rate lower than BA.

**Figure 6.**
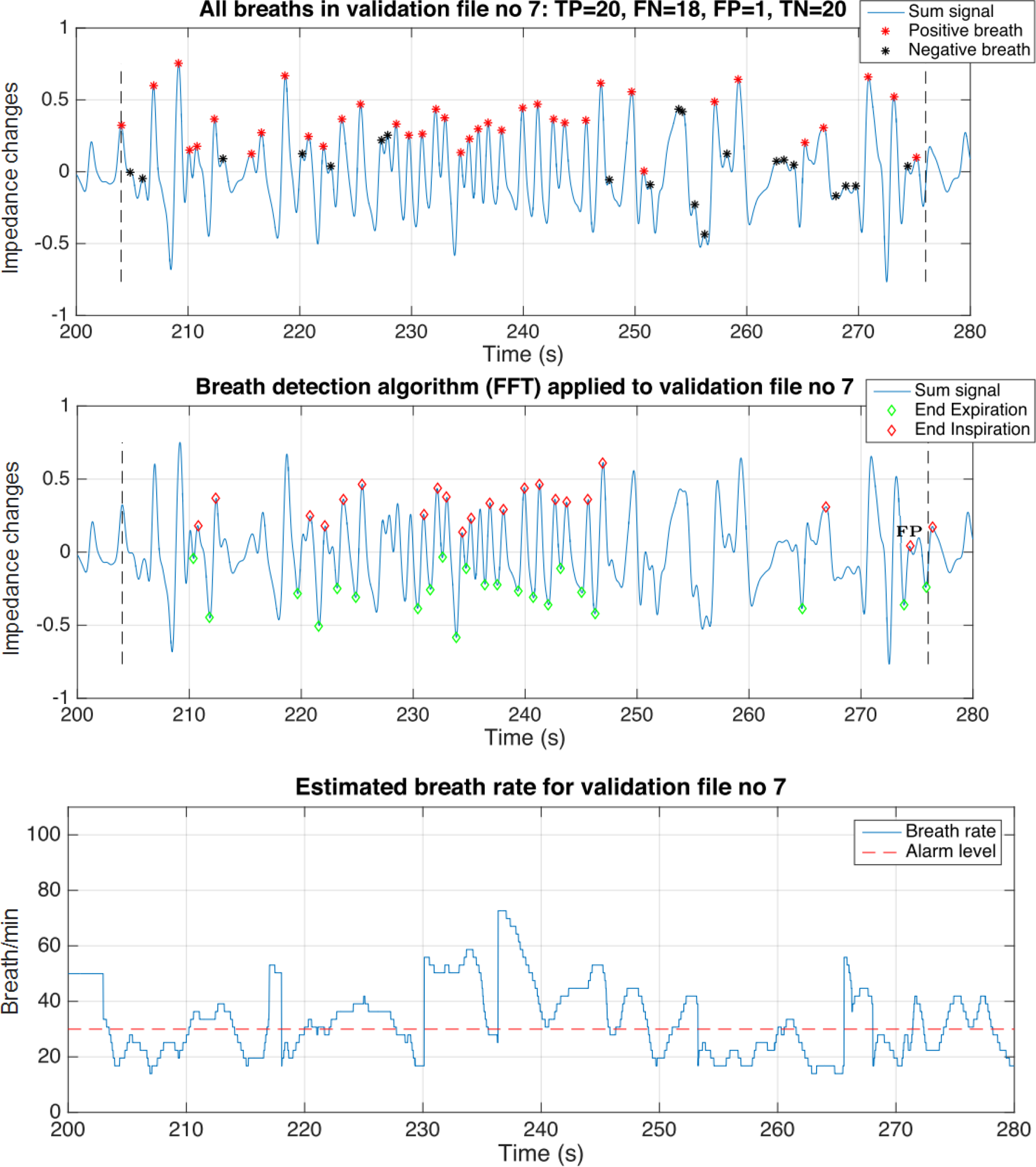
Breath detection performance on the validation data no.7 with zero-crossing algorithm complemented with amplitude threshold and FFT-based breath-rate estimation (ZC-AT-FFT). FP represents false positive.

The results for both test and validation data are illustrated in the corresponding Receiver Operating Characteristics (ROC) plots shown in Figure 7. The upper plot shows the results with the ZC-AT-FFT algorithm and the lower plot with the ZC-AT algorithm. The green circles indicate the performances obtained with the various parameter settings calculated over the 10 testing (training) data, the red asterisks indicate the optimal points over the test set and the blue asterisks the corresponding performances calculated over the validation set. The black asterisks and circles indicate the validated performance of the ZC algorithm, with default parameters and optimal parameters as described above, respectively. It is seen that both the ZC-AT-FFT and the ZC-AT algorithms perform better than both the default ZC and the optimised ZC algorithms in test and validation data set. With the current weight (w=10), the ZC-AT-FFT results in slightly lower FPR than the ZC-AT whereas, the ZC-AT provides slightly higher TPR than the ZC-AT-FFT algorithm.

**Figure 7.**
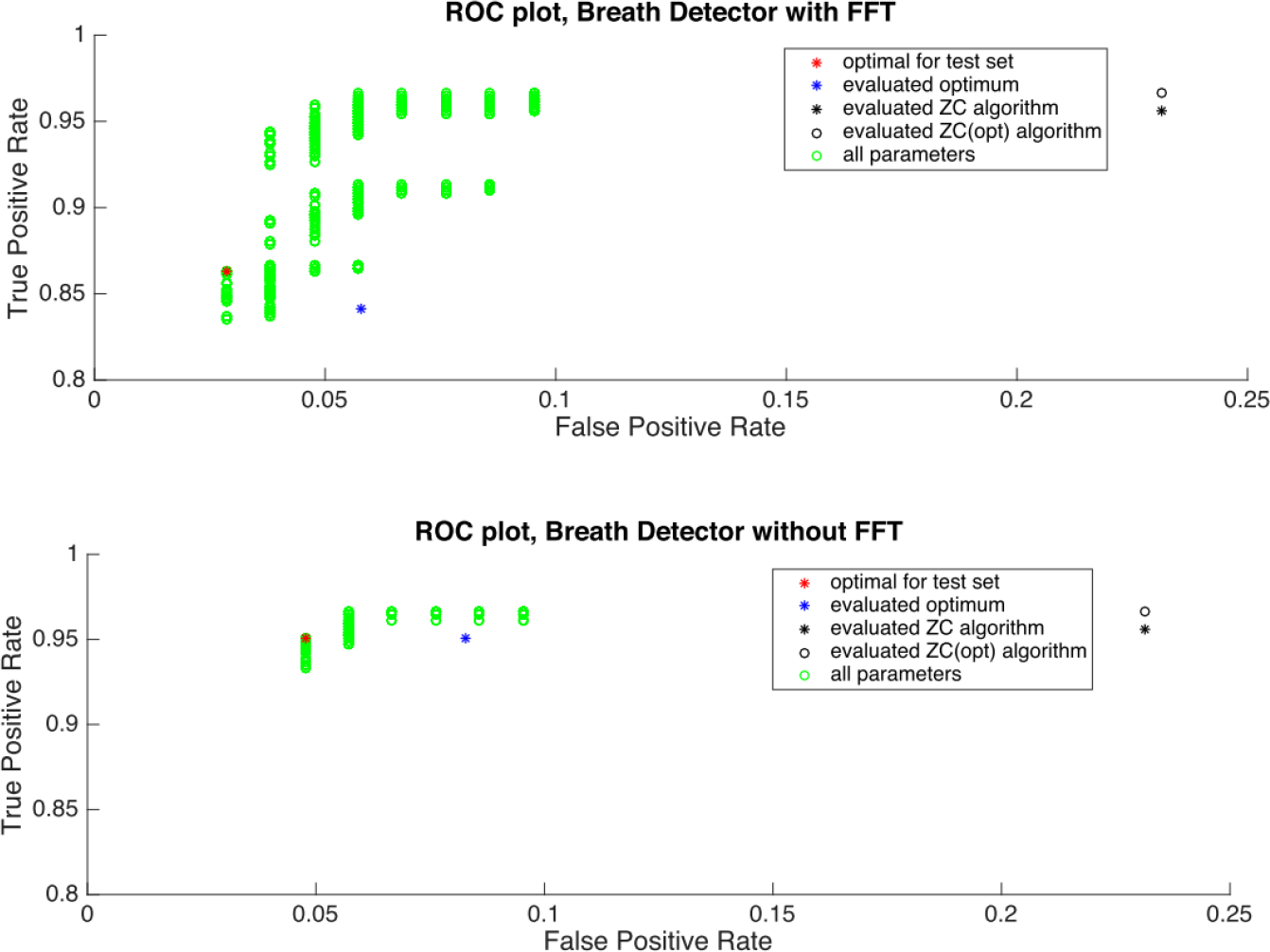
Receiver Operation Characteristics (ROC) for breath detection with the ZC-AT-FFT (upper plot) and ZC-AT (lower plot) algorithms, and a comparison with the ZC algorithm.

## 5. Summary and conclusions

A generic framework for optimizing the breath delineation algorithms used in EIT has been given in this paper. In particular, the approach is based on the definition of a gold standard for breath detection, the associated sensitivity and specificity measures, an adequate optimisation criterion defined on the Receiver Operating Characteristics (ROC) plane and a set of detector parameters to be optimised. Three different algorithms are proposed that are improving the breath detector performance by adding conditions on 1) maximum tidal breath rate obtained from zero-crossings of the EIT breathing signal (ZC algorithm), 2) minimum tidal impedance amplitude (ZC-AT algorithm) and 3) minimum tidal breath rate obtained from Time-Frequency (TF) analysis (ZC-AT-FFT algorithm).

The results show that both the ZC-AT and the ZC-AT-FFT algorithms outperform the conventional ZC algorithm in terms of the ROC. The main reason for this is that the amplitude threshold is able to avoid small amplitude disturbances (such as cardiac related signal components, etc) during periods of low or no breathing activity being interpreted as breaths. The addition of the FFT-based breath rate estimate is able to further decrease the FPR, but only at the expense of a decreased sensitivity. The ZC-AT-FFT algorithm has the advantage to output the auxiliary instantaneous, short-time estimate of the breath rate. Its disadvantage, however, is the higher computational complexity (of the short-time FFT) as compared to the ZC-AT algorithm.

## Acknowledgments

We acknowledge the funding from the European Union’s Framework program for research and innovation Horizon 2020 (CRADL, Grant No. 668259). The study is registered in a clinical trials registry (ClinicalTrials.gov, NCT02962505). It was approved by the ethics committees at three clinical study sites: the Emma Children’s Hospital, Amsterdam, Netherlands (Ethics number: METC 2016/184), the Arch. Makarios III Hospital, Nicosia, Cyprus (Ethics number: EEBK/EP/2016/32) and the Oulu University Hospital, Oulu, Finland (Ethics number: EETTMK 35/2017).

